# SMAD4 MH2 Mutations Disrupt CREBBP/EP300 Recruitment and TGF-β-Induced Transcription in Colorectal Cancer

**DOI:** 10.64898/2026.06.30.735541

**Authors:** Md Saiful Islam, Sheikh Nizamuddin, Timothy En Haw Chan, Omid Fotouhi, Stefanie Koidl, H.T. Marc Timmers

**Affiliations:** German Cancer Consortium (DKTK) partner site Freiburg, German Cancer Research Center (DKFZ), Germany and Department of Urology, Medical Center-University of Freiburg, Germany; International Max Planck Research School for Immunobiology, Epigenetics & Metabolism (IMPRS-IEM), Max Planck Institute of Immunobiology and Epigenetics, Germany

**Keywords:** SMAD4, colorectal carcinoma, TGF-β, missense mutation, transcription co-activators, gene transcription

## Abstract

SMAD4 is a central transcriptional effector of the TGF-β signaling pathway and a frequently inactivated tumor suppressor gene in various cancers. Missense mutations in its MH2 domain are among the most prevalent somatic alterations in colorectal cancer (CRC). These mutations are associated with disease progression and poor prognosis, yet their precise mechanistic consequences have remained incompletely characterized. Here, we show that CRC-derived SMAD4 MH2 hotspot mutations (D351H, S357P, R361C, and R361H) selectively impair co-activator recruitment without disrupting chromatin occupancy. RNA-seq profiling demonstrated broad suppression of TGF-β target gene expression across all mutants. Notably, the mutations confer distinct degrees of TGF-β pathway unresponsiveness: R361H is completely refractory to TGF-β stimulation, whereas R361C and S357P retain partial transcriptional responsiveness suggesting allele-specific differences in the severity of co-activator interface disruption. Genome-wide chromatin binding analysis by greenCUT&RUN confirmed that all mutants maintain wild-type-like genomic occupancy, as expected given that the MH1 DNA-binding domain is intact in each case. Proximity-dependent biotinylation mass spectrometry in COLO205 cells revealed that all four mutants exhibit markedly reduced interactions with the CREBBP/EP300 histone acetyltransferase complex and BRD4 relative to wild-type SMAD4 identifying disrupted co-activator engagement. Collectively, our findings establish that SMAD4 MH2 mutations impair TGF-β-induced transcription by selectively reducing CREBBP/EP300 recruitment, which provides a molecular mechanism for the loss-of-function SMAD4 phenotype in CRC.

**Graphical Abstract:** 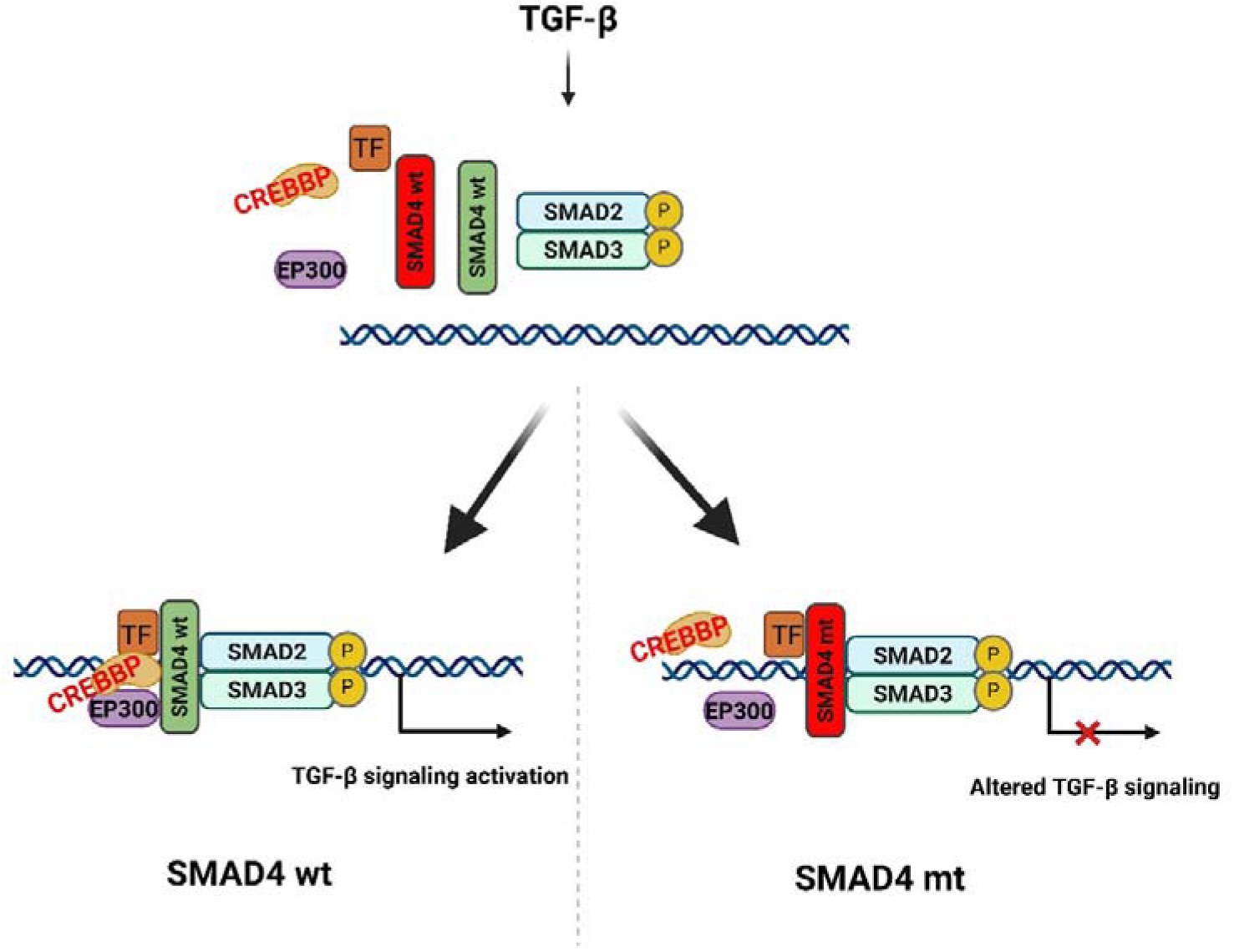

## Introduction

TGF-β signaling is a fundamental and versatile pathway in metazoans, regulating embryonic development, tissue differentiation, and homeostasis, while its dysregulation can lead to different human diseases.^1,2^ This signaling pathway becomes active, when TGF-β ligands bind type II receptors (TGFBRII), which cross activate type I receptors (TGFBRI). Subsequently, the activated receptor acts as a serine-threonine kinase to phosphorylate downstream effectors.^3^ Canonical signaling proceeds through phosphorylation of the cytoplasmic SMAD2 and SMAD3 transcription factors, which form a protein complex with SMAD4. The trimeric complex translocates to the nucleus^4,5,6^ in order to regulate transcription of TGF-β target genes in cooperation with transcription factors and co-activators.^7,8^ SMAD proteins are categorized into three groups: receptor-activated R-SMADs (SMAD1, SMAD2, SMAD3, SMAD5, and SMAD8), the common mediator Co-SMAD4, and inhibitory SMADs (SMAD6, SMAD7). Most of the SMAD proteins share structural similarity with two domains, MH1 and MH2.^8,9^ The MH1 domain interacts with DNA sequences, while the MH2 domain is responsible for protein-protein interactions including interactions with other SMADs, transcription factors, transcription co-factors, and receptors.^10,11^

TGF-β signaling exerts a tumor-suppressive role by inhibiting cell cycle progression, promoting apoptosis and preventing cell immortalization.^12^ However, gene mutations in TGF-β pathway components (*TGFBRI/II, SMAD2, SMAD3, SMAD4*) are frequently observed in cancers including colorectal cancer (CRC).^13^ In particular, *TGFBRII* is often mutated in microsatellite-instable CRC (10 -15% of cases), while *SMAD4* mutations occur in 10-30% of CRCs. *SMAD4* mutations occur predominantly as missense mutations as well as frameshift mutation, which result in expression of mutant SMAD4, reduced or complete loss of protein expression. CRC patients with reduced SMAD4 expression demonstrate significantly lower overall survival and reduced sensitivity to chemotherapeutic agents such as 5-fluorouracil.^14–16^. Nearly 90% of *SMAD4* mutations are missense in CRC, which primarily affects the MH1 and MH2 domains. MH1 mutations impair DNA binding and protein stability, while MH2 mutations disrupt interactions with SMAD2/SMAD3 and other partners proteins. *SMAD4* missense mutations are usually considered as loss-of-function mutations.^17,18^ Previous studies showed that SMAD4 MH2 domain mutations failed to activate TGF-β-induced gene expression^19^, which contributes to the aggressiveness of CRC. The molecular mechanisms behind this defective transcriptional activation are yet to explore.

In this study, we focus on a comparative analysis of CRC-derived *SMAD4* mutants with wild-type (wt) SMAD4. We ectopically expressed SMAD4 proteins tagged with GFP or miniTurbo using a doxycycline induction system in null-SMAD4 COLO205 cells to compare their functions at the transcriptomic, genomic and proteomic levels. Our results revealed that all SMAD4 mutants are defective in the transcriptional response to TGF-β treatment, which may result from reduced interactions with critical co-activator molecules like CREBBP/EP300.

## Materials and methods

### Cell culture and cloning

The human COLO205 (RRID: CVCL_F402) colorectal cancer cell line (purchased from ATCC, USA) and human retinal pigmented epithelial-1 (RPE-1; RRID: CVCL_4388) cell line were grown in a standard growth medium (DMEM, 10% FBS and 1% penicillin/streptomycin). *SMAD4* mutants were generated by site-directed mutagenesis PCR reaction using the *SMAD4* wild-type cDNA.^20^ A lentiviral plasmid vector pCW57.1 (Addgene #41393) was modified by inserting GFP or miniTurbo DNA sequences upstream of the attR1 site to create N-terminally tagged SMAD proteins. *SMAD4* wt and mutant cDNA constructs were transferred to the modified lentiviral vector downstream of GFP or miniTurbo sequences using the GATEWAY cloning technique (Thermo Fisher Scientific). All the plasmid constructs were confirmed by Sanger sequencing. GFP/miniTurbo-tagged SMAD4 wild-type and mutant cell lines were generated using the lentiviral transduction protocol. Transduced cells were selected using 1 µg/ml puromycin (Invitrogen, USA). All cell lines were confirmed as mycoplasma-free before performing the experiments.

### Cell treatment

Cells were treated with 1 µg/ml doxycycline (Thermo Fisher Scientific # J60579.22) overnight as indicated. Cells were treated with 10 ng/ml TGF-β (ProteinTech # HZ-1011), and an equal volume of DMSO as a control. The induction time of TGF-β and DMSO varied from 0 hours to 30 hours based on the experiment. For the CREBBP/EP300 experiment, cells were treated with 5 μM A-485 (Med Chem Express) for 4 hours prior to TGF-β treatment, and an equal volume of DMSO as a control.

### Immunoblotting

2 X Laemmli buffer (4% SDS, 20% Glycerol, 120 mM Tris-HCl pH 6.8) was used to prepare total cell lysates. Protein concentrations were measured using the BSA standard and Bradford assay (Sigma-Aldrich). The cell lysate was denatured at 95°C for 10 min and 50 µg protein were loaded per lane on 8% or 10% polyacrylamide/SDS gels. Proteins were transferred to PVDF membrane (Invitrogen) by liquid transfer protocol. Membranes were blocked for one hour at RT in TBS-T or PBS-T either with 5% milk or 5% milk/0.5% BSA, depending on the primary antibody (**Supplementary information**). According to the recommendations from the supplier, the primary antibody was diluted either in blocking buffer, and membranes were incubated with the primary antibody at 4°C for overnight. On the next day, membranes were washed three times (5 min intervals) with PBS-T or TBS-T buffer and incubated with a secondary antibody for 1 hour at RT. Blots were developed for imaging with the Clarity western ECL kit (Bio-Rad, Hercules, CA) and were imaged using a Chemidoc imager (Bio-Rad, Hercules, CA).

### qRT-PCR

Cellular RNA was isolated from cell lysates using RNeasy mini-kits (Qiagen). 1 µg of RNA was converted to cDNA using random hexamer primer and Superscript III reverse transcriptase according to the manufacturer protocol (Thermo Fisher Scientific). Following the manufacturer’s instructions, qRT-PCR was carried out in a CFX384 PCR instrument (Bio-Rad) with 10 ng of cDNA per reaction using these primer sets (**Supplementary Information**). The expression values of target genes were calculated using the 2^-ΔΔCt^ method and always normalized to the housekeeping genes (*GAPDH, TBP, ACTB, HPRT1, HBMS*). The expression values were presented as fold-change compared to the control samples unless otherwise stated.

### RNA-Seq

RNA was isolated from cell lysate using RNeasy mini-kits (Qiagen). The integrity and concentration of RNA samples were measured by a 2100 bioanalyzer using RNA 6000 Nano chips (both Agilent, USA) and Qubit 4 Fluorometer (Invitrogen, USA), respectively, in accordance with the manufacturer’s instructions. Only samples with a RIN value of 9.0 or higher were considered for the RNA sequencing. Genomic and proteomics core facility (DKFZ, Heidelberg) prepared the sequencing libraries from total RNA sample and used Illumina TruSeq standard total RNA library prep kit following the manufacturer’s instruction. Multiplex libraries were sequenced in a paired ending 100 bp setting using the NovaSeq 6000 instrument (Illumina, USA). The NGS data have been deposited with the accession number PRJNA965939 in the NCBI BioProject database (https://www.ncbi.nlm.nih.gov/bioproject/).

### Bioinformatics analysis of transcriptome datasets

Raw sequence data files were processed using snakePipes (v 2.4.1) ^21^ in mRNAseq mode and the generated feature count files were used for further analysis. The PCA was performed using DESeq2. The heatmaps were generated using Galaxy server.^22^ The pathway analysis was performed using clusterProfiler (version 4.4.3).

### greenCUT&RUN

To analyze the genome-wide binding profile of wt and mutant SMAD4 proteins, the greenCUT&RUN technique was applied to COLO205 cells expressing GFP-tagged SMAD4 wt and mutant proteins. In all cases, one million cells were harvested and washed with cold PBS. The cells were immobilized on concanavalin A-conjugated paramagnetic beads, permeabilized with 0.05% digitonin, and subjected to greenCUT&RUN protocol as described earlier.^23,24^ Mononucleosomal DNA from *Drosophila* S2 cells (20 pg) was added as spike-in DNA for normalization purposes. DNA libraries for sequencing were prepared using NEB Ultra II kits (New England Biolabs, USA). DNA quantity and size distribution of sequencing libraries were measured with the Qubit 4 fluorometer (Invitrogen, USA) and the 2100 Bioanalyzer (Agilent, USA) using DNA high sensitivity assay chips, respectively. DNA libraries were sequenced by the Max Planck Deep sequencing core facility. The NGS data have been deposited with accession number PRJNA965939 in the NCBI BioProject database (https://www.ncbi.nlm.nih.gov/bioproject/).

### Bioinformatics analysis of greenCUT&RUN data

100 bp paired-end sequencing reads were analyzed with 4-12 M reads per sample, respectively. The raw data were passed for quality control using Trim-galore (Version 0.6.4) with default parameters. The sequence reads that passed QC were aligned on the human (version of hg38) and Drosophila reference genome (BDGP5 release 75) using bowtie2 (version 2.3.5.1) with option: -dovetail -local -very-sensitive-local -no-unal -no-mixed -no-discordant -I 10 -X 700.^23,25^ An equal number of reads was adjusted for all samples, including SMAD4 wt and mutants, using Sambamba (version 0.8.0), and considered for further analysis. Homer was used for calling both narrow and broad peaks with default parameters except filtering based on clonal signals was disabled with option: -C 0. Homer was also used to annotate and find differential the peaks. The heatmaps were generated using computeMatrix function of the deeptools (version 3.5.1) with default parameters with an exception for the sum of the reads that were calculated per bin using the option - “averageTypeBins” instead of mean. To normalize the matrix sum, the reads were divided by total SpikeIn reads. PCA analysis was performed using DESeq2 package of R and pathway analysis was performed using clusterProfiler (version 4.4.3).

### Mass spectrometry analysis

COLO205 cells expressing miniTurbo-tagged SMAD4 proteins were grown in 10 15-cm dishes and expression was induced overnight with 1 µg/ml doxycycline. On the next day, cells were treated with 10 ng/ml TGF-β for 4 hours and the cells were incubated with 50 µM biotin in normal growth medium for 4 hours. To harvest the cells, media were aspirated and washed with cold PBS and cold buffer A (10 mM Hepes-KOH pH 7.9, 1.5 mM MgCl_2_, 10 mM KCl) sequentially. After aspirating this buffer, around 800 µl of complete buffer A (10 mM Hepes-KOH pH 7.9, 1.5 mM MgCl_2_, 10 mM KCl and freshly added 1X Roche CPI, 1 mM DTT, 0.2% NP40) were added on top of the dish and incubated on the ice for 5 min. Cells were harvested by cell scrapers and cytoplasmic fractions were collected after 6 mins of centrifugation at 3,200g and 4°C. Cell pellets were washed with buffer A to remove the remaining cytoplasmic protein. Nuclear pellets were resuspended in 2.5 ml of complete buffer C (420 mM NaCl, 20 mM Hepes-KOH pH 7.9, 20% v/v glycerol, 2 mM MgCl_2_, 0.2 mM EDTA and 0.2% NP40 and freshly added 1X Roche CPI, 1 mM DTT, 0.2% NP40) and agitated for 60-90 mins at 4°C. Finally, both the cytoplasmic and nuclear extracts were centrifuged at 40,000 *g* for 30 min at 4°C. The supernatants were collected for protein concentration measurement by the Bradford assay (BioRad, USA) and stored at -80°C until further use.

Streptavidin-beads (Cytiva, USA) were washed two times with a binding buffer (20 mM Hepes-KOH pH 7.9, 420 mM NaCl, 20% v/v glycerol, 2 mM MgCl_2_, 0.2 mM EDTA, 0.2% NP-40 with freshly added 1X Roche CPI and 1 mM DTT). For control pulldown, half of the beads was incubated with 1 mM biotin in binding buffer and the other half was incubated with binding buffer for 60 min at 4°C. Beads were washed three times with binding buffer. 1 mg of nuclear protein lysate was added to the washed beads per pulldown, which was performed in triplicate for both the streptavidin pulldown and the biotin-blocked streptavidin control pulldowns. The pulldown reactions were incubated rotating for 60-75 min at 4°C. The beads were washed twice with binding buffer, twice with PBS with freshly added 1X Roche CPI, 1 mM DTT and 0.2% NP-40) and twice with plain PBS.^26^ On-bead digestion of bound proteins was performed in elution buffer (100 mM Tris-HCl pH 7.5, 2 M urea, 10 mM DTT) with 0.1 µg/ml trypsin overnight at 22°C. TFA was added to 1% after, which tryptic peptides were bound to C18 stage tips (ThermoFischer). USA. Tryptic peptides were eluted in with 80% acetonitrile and dried by lyophilization.

Lyophilized tryptic peptides were suspended in 10% formic acid prior to MS analysis and one-third of resuspended samples were analyzed as described in^27^ with minor modifications. Nanoflow LC-MS/MS analyses were carried out using an Easy nano-LC 1200 system (Thermo Fisher Scientific) interfaced with an Orbitrap Fusion Lumos mass spectrometer. Peptides were resolved on a 25 cm C18 reversed-phase column (75 μm internal diameter, 2 μm particle size, 100 Å pore size), using a binary gradient of increasing organic content (mobile phase A: 0.1% formic acid in water; mobile phase B: 0.1% formic acid in 80% acetonitrile) delivered at a flow rate of 300 nl/min. Tandem mass spectra were acquired using a data-dependent TOP-10 acquisition method. After a single MS scan, a maximum of 10 MS/MS scans were performed in data-dependent mode. To prevent sample carry-over, blank samples of 10% formic acid were run for 45 minutes between the streptavidin pulldown and the biotin-blocked streptavidin control pulldown.

The raw data files were analyzed with MaxQuant software (version 1.5.3.30) using the Uniprot human FASTA database^28^ for alignment. Label-free quantification (LFQ) values and matches between run options were selected. Additionally, the intensity-based absolute quantification (iBAQ) method was activated to calculate the subsequent relative protein abundance. MaxQuant derived txt files were further processed by Perseus software (MQ package, version 1.6.12) to generate a volcano plot. In the Perseus analysis, contaminants and reverse hits were filtered out.^28^ LFQ intensity-based values were transformed on the logarithmic scale (Log2) to generate Gaussian distribution of data that allows for the imputation of missing values on normally distributed data (width =0.3, shift = 1.8). Protein enrichment between streptavidin pulldown and the biotin-blocked streptavidin control pulldown was calculated using a two-tailed student’s t-test, considering a false discovery rate (FDR) of 5%. The constant value of 1 was considered as the threshold of significance (S0=1). iBAQ values were used to determine the relative stoichiometry of selective interactors.^29,30^ All proteomic data are available from the PRIDE database (PXD041662).

## Results

### Ectopic expression of wild-type and mutant SMAD4 proteins in COLO205 cells

The aim of our study is to examine the effect of cancer mutations on the function of SMAD4 in TGF-β signaling. In order to select CRC-derived *SMAD4* mutations we examined the COSMIC and The Cancer Genome Atlas (TCGA) databases. These database searches revealed a *SMAD4* mutation frequency of 12.6% in CRC samples. A total of 652 mutations were detected out of 4,398 patient samples. Among these mutations, 478 were identified as putative driver mutations, which are known or predicted to confer a selective growth advantage to cancerous cells. The remaining 174 mutations are known as passenger mutation with unknown significant role in tumorigenesis. A majority of driver mutations are missense (68%, 329 of 478), while truncating mutations account for 26%. The rest of the driver mutations include a very small percentage of in-frame, splicing and fusion mutations. A large proportion of the *SMAD4* missense mutations (288/652) reside in the MH2 domain (**Fig. S1A, S1B**).

For this study we selected four MH2 mutations, which frequently occur in CRC. As a first step we examined whether these missense mutations affect protein stability or expression. For this, we selected the CRC cell line COLO205, which lacks SMAD4 protein expression due to homozygous deletion of the first four exons of *SMAD4*.^31^ Lentiviral transduction was employed to generate COLO205 cells expressing dox-inducible GFP-tagged versions of wild-type (wt) and SMAD4 mutant proteins. Immunoblotting was performed with whole cell lysates of the transduced cell lines using antibodies directed against GFP or SMAD4, which confirmed expression of wt and mutant SMAD4 proteins (**Fig. 1A**). In addition to the CRC cell line, we studied the expression of those mutant SMAD4 proteins in a retinal pigmented epithelial cell line (RPE-1) using lentiviral transduction. Immunoblot analysis showed similar expression levels of GFP-tagged SMAD4 wt and mutants, which were higher than endogenous SMAD4 (**Fig. S1C**). Similar levels GFP-tagged SMAD4 proteins indicated that these MH2 missense mutants are not affecting protein expression or stability.

**Figure 1:**
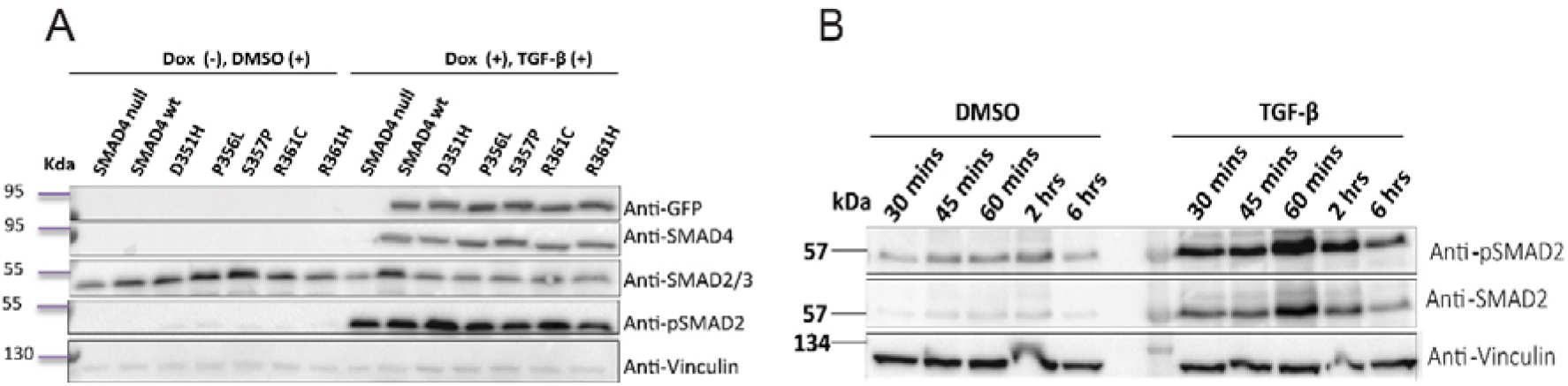
Ectopic expression of wild-type and mutant SMAD4 proteins. A) Immunoblot analysis of GFP-tagged SMAD4 proteins expressed in COLO205 cells, B) Immunoblot analysis for phosphorylated SMAD2 (pSMAD2) upon TGF-β treatment. Cells were treated with 10 ng/ml TGF-β for the respective time points and an equal volume of DMSO was used as a control.

### SMAD4 MH2 domain mutants failed to activate TGF-**β** induced genes

To investigate activation of the TGF-β signaling pathway in COLO205 cells, we performed immunoblotting analysis to examine phosphorylation of SMAD2 as a read-out. COLO205 cells expressing wt SMAD4 protein were treated with 10 ng/ml TGF-β in a time series experiment to determine the peak of SMAD2 phosphorylation. Immunoblot analysis indicated induction of phosphorylated SMAD2 (pSMAD2) already 30 mins after TGF-β addition, which gradually increased to its peak at 60 min (**Fig. 1B**). Based on pSMAD2 analysis, null SMAD4, wt SMAD4 and mutant SMAD4 cells were treated with 10 ng/ml TGF-β for one hour before protein isolation. Immunoblot analysis using GFP and SMAD4 antibodies showed similar GFP-SMAD4 expression levels in all dox-induced cells, except and as expected for null-SMAD4 cells (**Fig. 1A**). And also as expected, endogenous SMAD2 expression was observed in all cell lines, which is independent of GFP-SMAD4 expression. TGF-β treatment induced phosphorylation of endogenous SMAD2 confirming that the upstream part of the TGF-β pathway is intact in COLO205 cells (**Fig. 1A**).

Next, we investigated the genome-wide TGF-β transcription signatures by RNA-seq of wt and mutant SMAD4 cells. We treated cells with TGF-β from 0 to 30 hours to perform RNA sequencing in triplicates for each cell line. PCA analysis of the top 500 genes showed that the null SMAD4 samples cluster separate from wt and mutant SMAD4 samples in principal component 1 (PC1) (**Fig. S2**). Upon TGF-β treatment, the wt SMAD4 time points cluster together in PC2, which is clearly different from the mutant SMAD4 clusters (**Fig. S2**). Taken together the PCA analysis indicates that the SMAD4 MH2 mutants are transcriptionally non-responsive to the TGF-β treatment.

3,200 differentially expressed (DE) genes were identified and categorized into four different clusters by unsupervised hierarchical clustering (**Fig. 2A**). Among them, cluster 3 genes are rapidly induced in wt SMAD4 but are only poorly induced in the SMAD4 mutants. The genes in cluster 4 are only induced at late timepoints and this is not affected by the SMAD4 mutations. Interestingly, cluster 1 and 4 genes remain flat in null SMAD4 cells. Cluster 2 genes are reduced in expression at late time points of TGF-β treatment in SMAD4 wt and mutant expressing cells and much less in SMAD4 null cells. Cluster 1 genes are also repressed at later time points, but this is not dependent on the SMAD pathway (**Fig. 2A**). Taken together, the time course RNA-seq experiment indicated that early induction genes are affected by the MH2 mutants of SMAD4 found in CRC, while late transcriptional effects of these mutants are much less pronounced.

**Figure 2:**
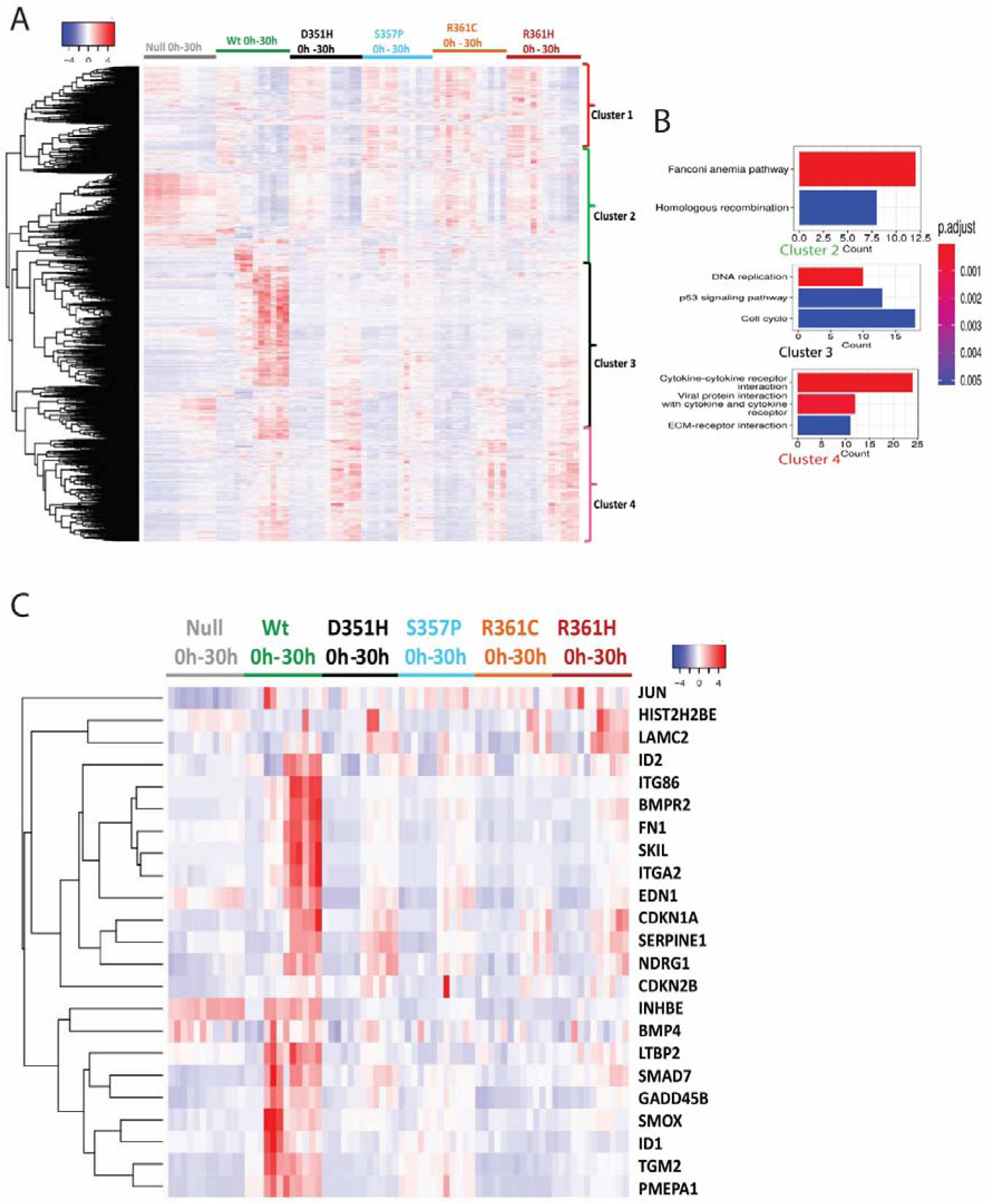
**TGF-**β **responsive genes of mutant and wt SMAD4 cells.** A) Heat map and clustering of differentially expressed genes in wt and mutant SMAD4 COLO205 cells upon TGF-β induction (0, 2, 18 and 30 hours), B) KEGG pathway analysis using cluster 2, 3 and 4 genes from RNA-seq clusters and C) Heat map of TGF-β core component genes. Cells were treated with 10 ng/ml TGF-β for the indicated times and equal volume of DMSO was used as control.

Next, we performed pathway analysis using the TGF-β responsive genes from different clusters. The KEGG pathway analysis revealed that wt SMAD4 upregulated genes residing in cluster 3 are involved in DNA replication, cell cycle and P53 pathway (**Fig. 2B**), while cluster 2 and 4 genes represent cytokine related pathways. Besides these, biological pathway analysis of cluster 3 genes showed several top hit pathways that are involved in cell cycle and DNA replication (**Fig. S3).** Surprisingly, we did not find the enrichment of direct TGF-β related pathways. In order to check the expression of core TGF-β responsive genes, we performed the heat map analysis of a list of genes, that are directly regulated by TGF-β signaling. The TGF-β core component genes were activated in wt SMAD4 expressing cells from early to late TGF-β induction time points. In contrast, these genes were non-responsive to TGF-β in cells expressing SMAD4 mutants and largely behaved like the parental COLO205 null SMAD4 cells (**Fig. 2C**). Taken together, the RNA-seq analysis of GFP-SMAD4 expressing COLO205 cells indicates that wt SMAD4 expression restores the transcriptional response of TGF-β target genes, while SMAD4 MH2 mutants strongly reduces the response to TGF-β.

### Mutant SMAD4 effectively binds with DNA

GFP-tagging of SMAD4 allows the genome-wide localization of non-crosslinked intact cells via the greenCUT&RUN approach. GreenCUT&RUN depends on micrococcal nuclease (MNase) fused to a GFP nanobodies and subsequent activation of nuclease activity by Ca^+2^ to release DNA-bound GFP-protein complexes.^24^ To investigate the genome-wide binding of wt and mutant SMAD4 proteins, we performed greenCUT&RUN experiment using our COLO205 cells expressing the GFP-SMAD4 proteins (wt, S357P, R361C, and R361H). To analyze the binding pattern of wt and mutant GFP-SMAD4, cells were treated with TGF-β in a time course experiment and libraries were prepared from released DNA and subjected to deep sequencing. The read numbers from each sample were normalized to a spike-in control. The data analysis identified a total number of peaks within a range of 12,000 to 35,000 in mutant and wt SMAD4 samples at different TGF-β treatment times (**Fig. S4A**). Upon TGF-β induction the coverage increased for wt and all mutants. Maximum coverage was obtained at 18 hours of TGF-β treatment in all samples. In total, ∼30% peaks contain the SMAD4 consensus motif (**Table S1**). Genome-wide binding site analyses indicated an almost similar pattern of binding between wt and mutant SMAD4 proteins to different genomic regions, except SMAD4 wt and R361C mutant, which display increased coverage at intergenic and intronic sites (data not shown). Genomic tracks of the TGF-β responsive *SERPINE1, PMEPA1,* and *ID1* genes showed binding either to their promoter and/or or upstream regions (**Fig. 3A, 3B, S4B**). Differential peak analysis revealed that as expected TGF-β treatment increased DNA binding of SMAD4. The early and late of TGF-β time points resulted in 830 (4.12%) and 1189 (6.16%) differential binding sites for wt SMAD4, respectively (**Fig. 3C**). Of these peaks 390 (46.9% of 830) and 549 (46.17% of 1189) peaks contained the consensus SMAD4 motif. The MH2 SMAD4 mutants behave largely similarly to wt SMAD4 confirming that these mutations do not affect the DNA binding potential of SMAD4, which is mediated by its MH1 domain.

**Figure 3:**
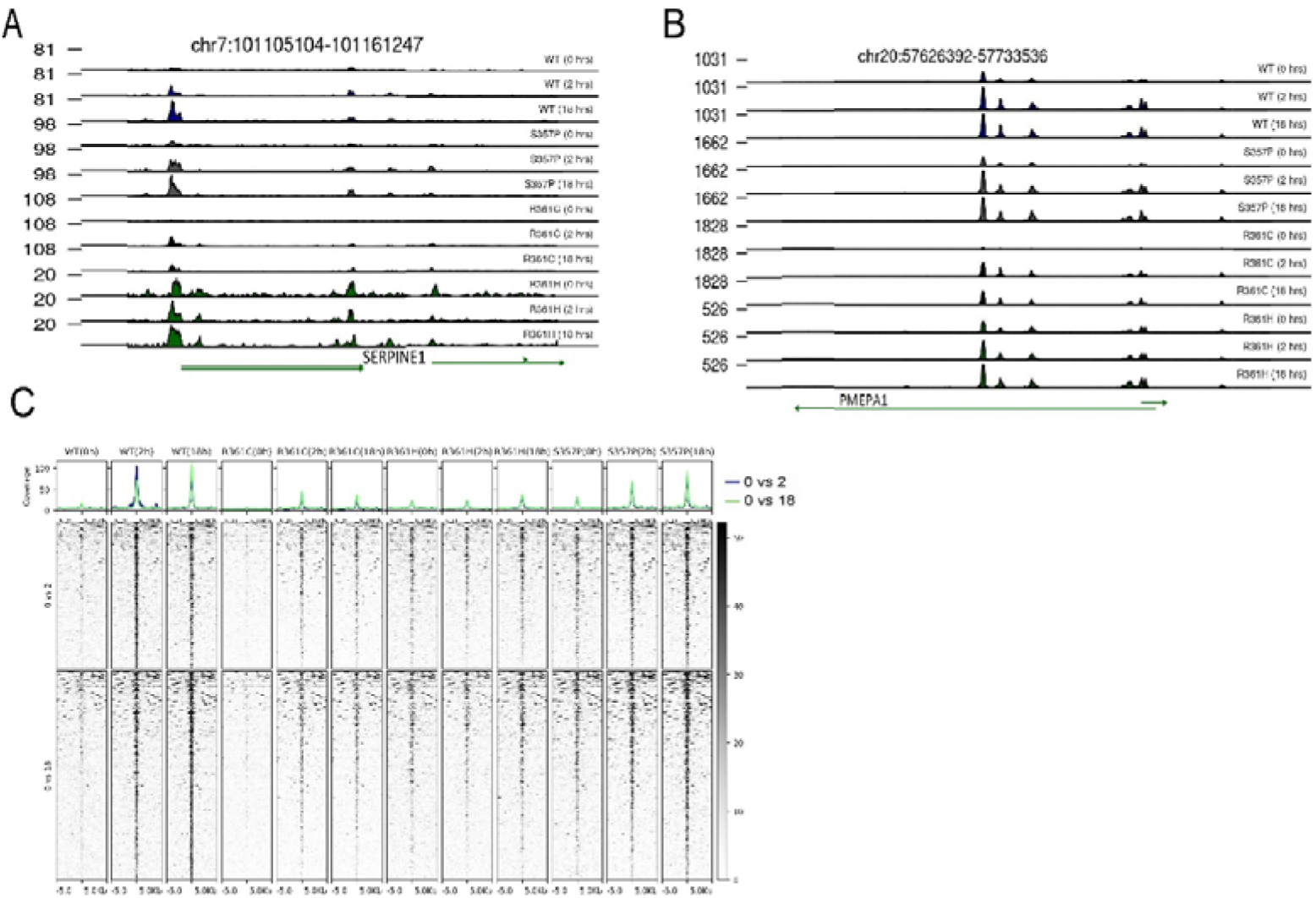
**Genome-wide binding analysis of wt and mutant SMAD4**. A) Coverage tracks of wt and mutant SMAD4 binding at the *SERPINE1* locus, B) Coverage tracks of SMAD4 proteins of the *PMEPA1* locus, C) Heat maps presenting genome-wide wt and mutant SMAD4 binding after TGF-β treatment.

Gene ontology analysis of the differential peaks did not reveal TGF-β associated enrichment for both the early and late induced genes. However, in molecular signature-based enrichment analysis (msigdb M2445), (FDR p-value = 6.9×10^-5^ and 7.5×10^-4^ for early and late TGF- β induction, respectively) and M2839 (FDR p-value = 1.8×10^-3^ and 3.1×10^-3^ for early and late TGF-β induction, respectively) gene sets were enriched (**Table S2**). The M2445 signature consists of genes, which were up-regulated in MEF cells (mouse embryonic fibroblast) upon stimulation with TGF-β for 10 hours, while M2839 consists of genes, which were up-regulated by TGF-β in a panel of epithelial cell lines. The coverage analysis around these 830 and 1189 differential peaks revealed that binding of SMAD4 mutants is also increased upon TGF-β induction (**Fig. 3C**).

Next, we correlated the transcriptomic data with genome-wide SMAD4 binding patterns upon TGF-β activation. The coverage analysis was focused at the promoter regions of the genes from different clusters. While we did not observe differences in coverage among the clusters for the same samples (**Fig. S4C**), the coverage increased in both early and late induction time. In conclusion, the analyzed CRC mutations in the MH2 domain of SMAD4 do not interfere with the genome binding of SMAD4.

### Mutant SMAD4 have reduced interaction with co-activators

As the SMAD4 MH2 domain mutations affect the transcription of TGF-β responsive genes, we hypothesized that MH2 domain mutations might be affecting SMAD4 interaction with other proteins. To systematically investigate the SMAD4 protein-protein interaction network in the context of TGF-β signaling, a proximity biotinylation labeling-based quantitative mass spectrometry (qMS) was set up using HA-miniTurbo system.^32^

For this purpose, dox-inducible miniTurbo-tagged SMAD4 wt and mutant protein expressing COLO205 cells were generated using lentiviral transduction. Comparable expression levels of all SMAD4 proteins were observed by immunoblotting analysis using HA epitope and SMAD4 antibodies (**Fig. S5A**). Nuclear extracts from TGF-β treated cells (10 ng/ml, 4 hours) with concomitant biotin labeling were subjected to streptavidin-based affinity purification and qMS. Enrichment of transcriptional co-activators including CREBBP, EP300, BRD4, KDM1A, and multiple MLL3/4 complex subunits upon TGF-β was observed in wt SMAD4 cells (**Fig. 4A, 5A**). As expected, SMAD2/SMAD3 proteins were not enriched in uninduced conditions, which validates specificity of this approach.

**Figure 4:**
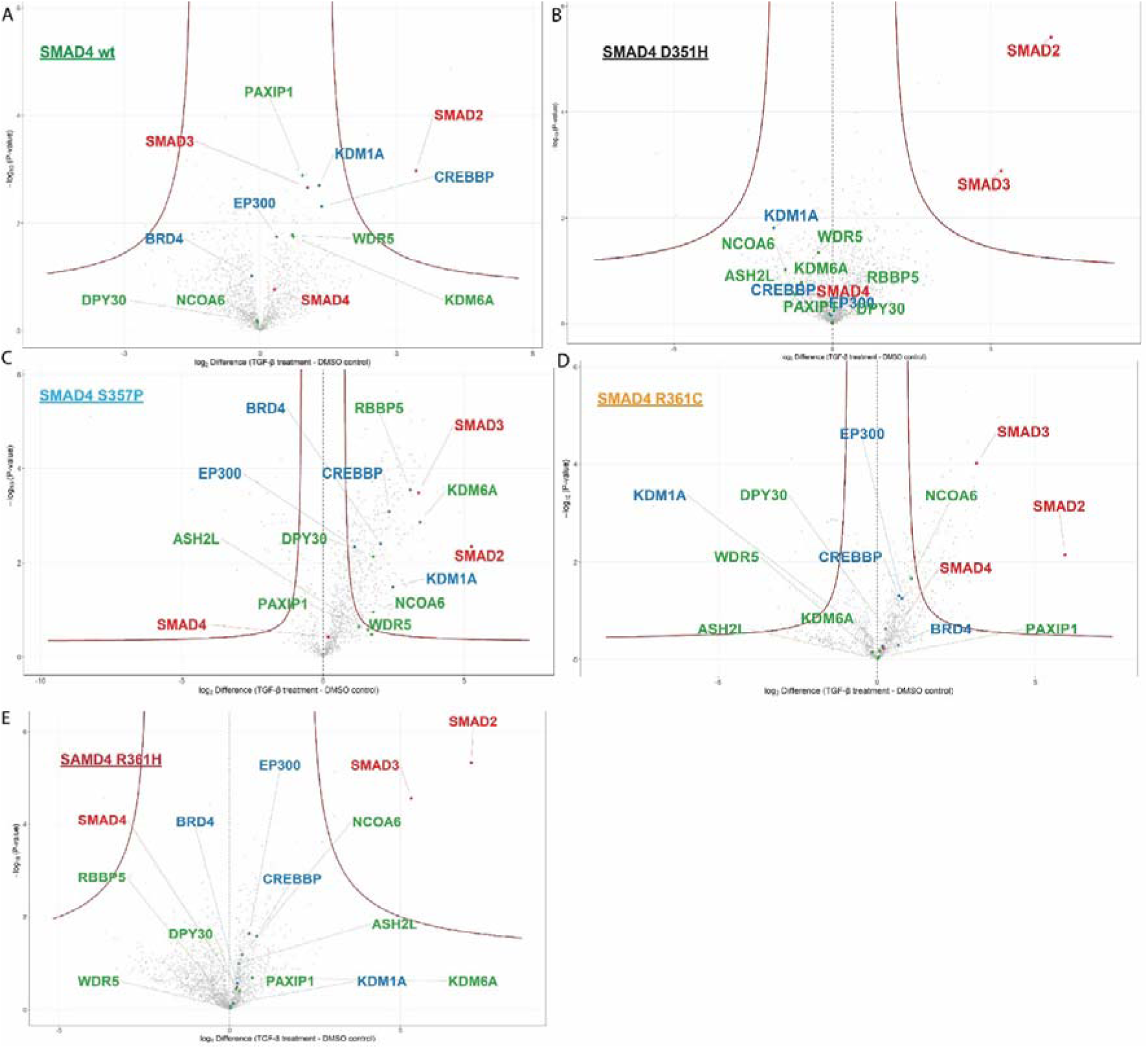
miniTurbo based proximity labeling followed by qMS for wt and mutant SMAD4. Volcano plot analysis for co-activator, MLL3/4 complex proteins in A) wild-type SMAD4, B) D351H, C) S357P, D) R361C, and E) R361H nuclear extracts. Cells were treated with 10 ng/ml TGF-β for 4 hours and 50 µM biotin. DMSO was used as a control treatment.

Next, comparative qMS analysis of the SMAD4 MH2 mutants (D351H, S357P, R361C and R361H) were performed to determine enrichment of co-activators, transcription co-factors, and subunits of MLL3/4 protein complex. This indicated that a reduction (S357P, R361C and R361H) or complete loss (D351H) of CREBBP, EP300 and KDM1A labeling (**Fig. 4B-E**, **Fig. 5B-E**). On the contrary, most of the subunits of MLL3/4 protein complex were enriched in the SMAD4 mutants like as SMAD4 wt except for D351H mutant (**Fig. S4B-F**).

**Figure 5:**
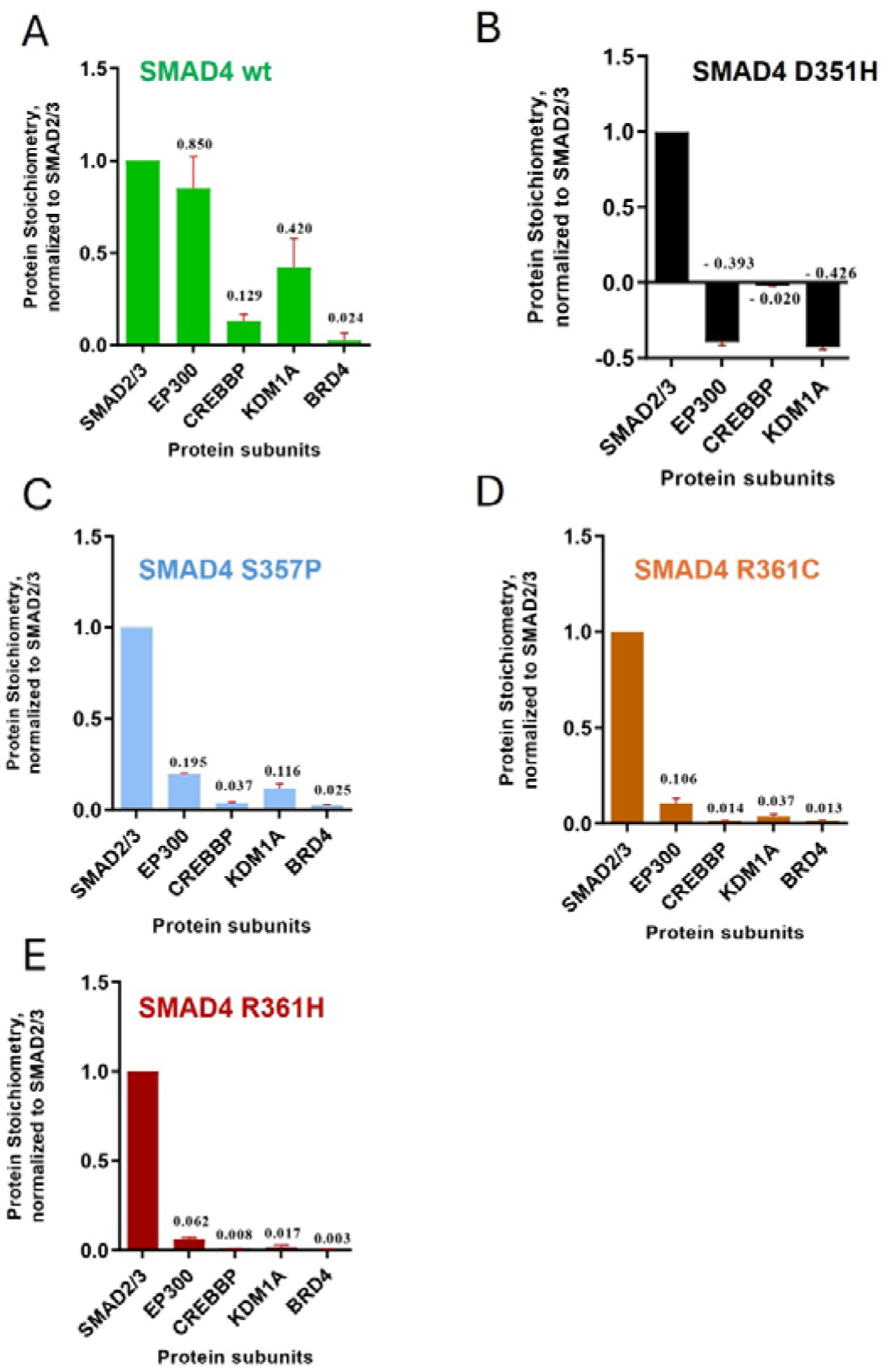
Co-activator enrichment of SMAD4 proteins using proximity-labeling by miniTurbo followed by quantitative mass spectrometry. Relative labeling of co-activator proteins in A) wild-type SMAD4, B) D351H, C) S357, D) R361C, and E) R361H nuclear extract. Cells were treated with 10 ng/ml TGF-β for 4 hours and 50 µM biotin.

These data indicate that compared to the wt SMAD4, the SMAD4 MH2 mutants displayed reduced interactions with co-activator proteins, most notably the CREB binding protein (CREBBP) and EP300, which provides a molecular mechanism for the transcriptional defects of MH2 mutations.

### CREBBP/EP300 is vital for the activation of TGF-**β** target gene

To investigate the roles of the CREBBP and EP300 co-activators in TGF-β-mediated gene activation, wt SMAD4-expressing COLO205 cells were treated with the CREBBP/EP300-inhibiting A-485 molecule, which is a competitive inhibitor of acetyl-CoA binding for the histone acetyltransferase activities (HAT) of CREBBP and EP300.^33^ Quantitative RT-PCR analysis revealed that A-485-treated cells exhibited significantly diminished the transcriptional responses of selected TGF-β target genes (**Fig. S6A**).

To comprehensively assess transcriptional changes, we performed RNA-seq analyses of wt SMAD4 cells treated with A-485. PCA analysis of the RNA-seq data (**Fig. S6B**) revealed distinct clustering of samples according to treatment and time, which validates data quality. Volcano plots of the RNA-seq data revealed that under basal conditions (without TGF-β treatment), that as expected A-485 treatment by itself caused alterations in gene expression compared to untreated cells (**Fig. 6A**). While a robust activation is observed in control cells for genes like *ITGA2, FERMT1, SKIL, ITGB6, JUN, INHBE, PMEPA1, LAMC2, SERPINE1* and *TGFB1*, this induction was reduced upon pretreatment with A-485 (**Fig. 6B)**. Hierarchical clustering and heatmap visualization confirmed the reduced transcriptional responses to TGF-β program following CREBBP/EP300 inhibition. The heatmap distinctly separated cells treated with TGF-β alone from those pre-treated with A-485 prior to stimulation (**Fig. 6C**), which revealed a strongly reduced induction of key TGF-β target genes such as *ID2, INHBE, SKIL, ITGA2, ITGB6* and *JUN* upon CREBBP/EP300 inhibition. Together, these results demonstrate that the HAT activities of the CREBBP and EP300 co-activators are indispensable for the transcriptional activation of TGF-β target genes. Loss of CREBP/EP300 function critically compromises this pathway, which supports the conclusion that transcriptional defects displayed by the CRC-derived SMAD4 MH2 mutations are related to loss of CREBP/EP300 interactions.

**Figure 6:**
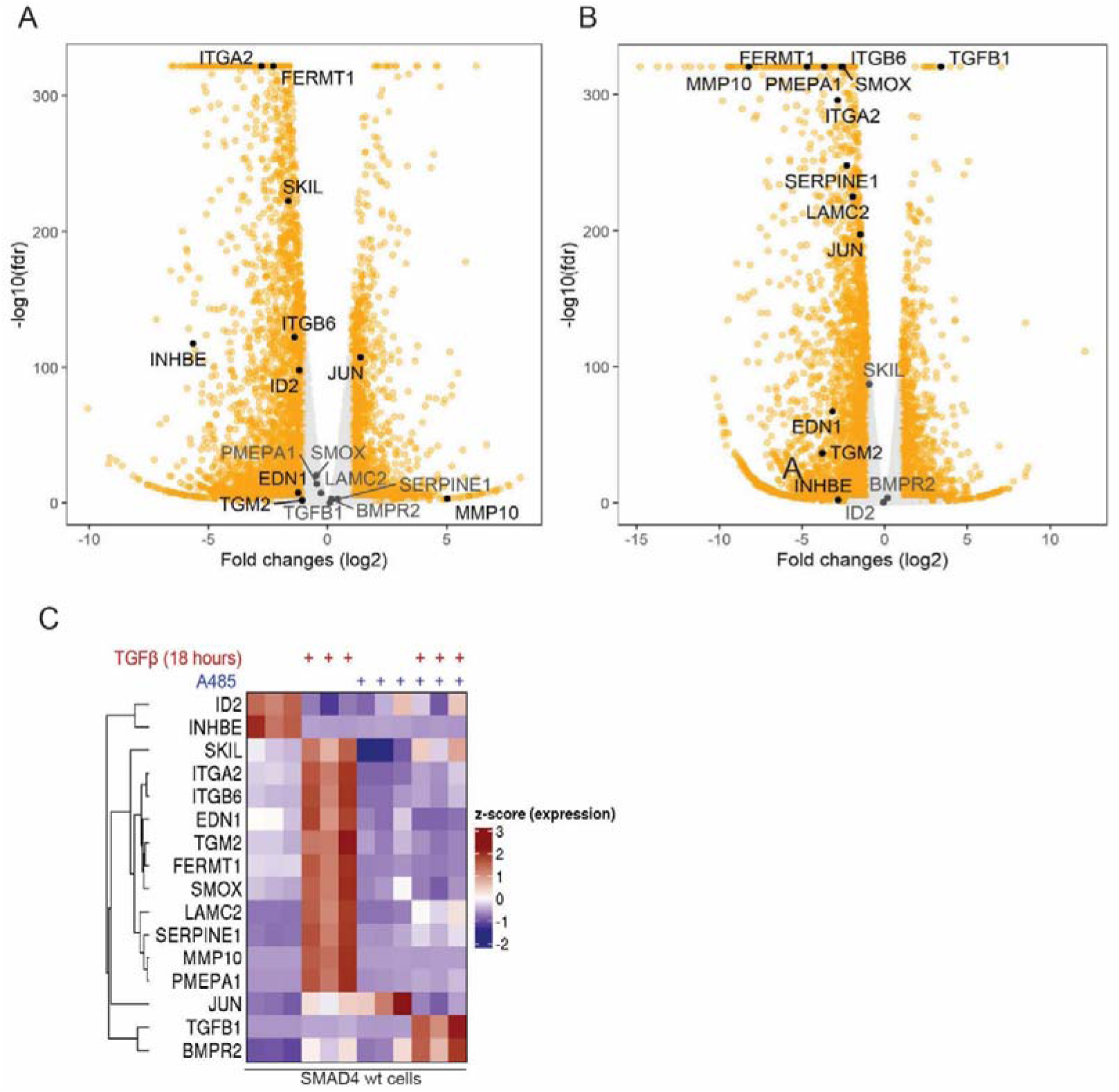
**Genome-wide TGF-**β **target gene expression upon CREBBP/EP300 degradation.** A) Volcano plot analysis to compare the differential gene expression in A-485 cells vs untreated cells without TGF-β induction, B) A-485 treated cells vs untreated cells followed by 18 hours of TGF-β induction, 2276 differentially expressed genes were considered. C) Heatmap analysis using the gene sets regulated by TGF-β signaling pathway. Cells were treated with 10 ng/ml TGF-β for 18 hours and A-485 treatment for 4 hours at 5 µM.

## Discussion

SMAD4 is a pivotal transcriptional effector in the TGF-β signaling pathway as well as one of the most frequently mutated tumor suppressors in colorectal cancer (CRC). While *SMAD4* mutations in CRC are generally considered as loss-of-functions, the precise molecular consequences of its MH2 domain missense mutations have remained poorly defined.^34,35^ A central and unresolved question has been whether MH2 mutations abrogate TGF-β target gene expression by interfering with ability of SMAD4 to bind to chromatin or, alternatively, by selectively disrupting its capacity to recruit transcriptional co-activators at target loci. Our study resolves this question and, in doing so, identifies a molecular vulnerability with potential clinical relevance.

In this study, we selected four frequently-occurring SMAD4 MH2 domain mutations (D351H, S357P, R361C and R361H) to study the role of these mutations in transcriptional responses to TGF-β. Comparative protein expression analysis of wild-type SMAD4 with the missense mutants revealed that the mutations are not interfering with stability or expression of the SMAD4 protein. Genome-wide gene expression analysis by RNA-seq showed that SMAD4 reintroduction supports the activation of well-known TGF-β responsive genes, while mutant SMAD4 expressing cells display a severely reduced response. As expected, genome-wide SMAD4 binding analysis showed that mutant SMAD4 proteins bind like their wt SMAD4 counterpart. SMAD4 recognizes SMAD binding element (SBE) motifs even at the non-induced state and treatment of TGF-β increases SMAD4 binding. Interestingly, the R361C mutant had lower binding under non-induced conditions compared to wt and the other SMAD4 MH2 mutants. Next, we correlated the RNA-seq data with genome-wide SMAD4 binding profile. Among the RNA-seq clusters (cluster 1-4), the genes of cluster 3 were strongly activated by TGF-β in wt SMAD4 cells. The wt and mutant SMAD4 occupancy did not change for RNA-seq gene cluster 3, although this cluster mostly contains genes activated by wt SMAD4. This indicates that the SMAD4 MH2 domain mutations are not hindering the interaction of the MH1 domain with DNA.^18^

In general, gene-specific transcription factors like the SMAD proteins interact with other transcription factors and co-activators to achieve efficient and proper transcription responses of their gene targets.^36^ Usually, gene-specific transcription factors recognize 4-8 bp long DNA sequences within the genome and their binding can be influenced by other gene-specific transcription factors, including AP-1 transcription factors for SMAD complexes. Many studies showed that gene-specific transcription factors can recruit co-activators to target DNA.^37^ These co-activators can relieve the repressive effect of nucleosomes and promote the transcription of specific genes by allowing the recruitment of the basal transcription machinery including RNA polymerase II (pol II) to gene promoters.^38,39^ Baas and coworkers identified the interaction of MLL4 co-activator complexes with the SMAD-specific transcription machinery to stimulate pol II-mediated transcription, ^20^ which involves a feed-forward loop of MLL4 with the AP-1 family member JUNB as we reported recently.^7^ Indeed, we identified several MLL4 complex members in the proximity-labeling qMS experiment with wt SMAD4 cells **(Fig. 4A).** We also observed enrichment of other transcription factors and co-activators was found upon TGF-β induction. Interestingly, proximity-labeling of uninduced samples detected most of those co-activators and transcription factors, while SMAD2/SMAD3 enrichment was observed only upon TGF-β treatment. This is in line with studies showing that the heterotrimeric SMAD2/SMAD3/SMAD4 complex requires active TGF-β signaling. ^40,41^ SMAD4 has both nuclear and cytoplasmic localizations in the uninduced state ^40,42^, which explains the detection of co-activators and transcription factors as SMAD4 interactors in the uninduced state. Importantly, MH2-mutant SMAD4 proteins interacted with SMAD2/SMAD3 after TGF-β treatment, which indicated that the SMAD4 mutants are not defective in formation of the SMAD2/SMAD3/SMAD4 trimer. In contrast to SMAD2 and SMAD3, the recruitment of co-activators following TGF-β induction was markedly reduced in cells expressing SMAD4 MH2 mutants. Most notably, enrichment of CREBBP/EP300 and BRD4 was diminished by approximately 3 to 20-fold compared to wt SMAD4. We propose that this impaired co-activator interaction underlies the defective TGF-β-dependent gene activation observed in the MH2 domain mutants in SMAD4.

To test the importance of the CREBBP/EP300 histone acetyltransferases for TGF-β activation in our cell system, we performed transcriptomic analyses using selective inhibitors to inactivate the CREBBP/EP300 HATs in COLO205 cells expressing wt SMAD4. Inhibition of CREBBP/EP300 abolished transcriptional activation of TGF-β target genes in wt-SMAD4 expressing COLO205 cells. These results are in line with previous studies demonstrating a critical role for CREBBP/EP300 in SMAD-dependent, TGF-β-induced gene expression in primary endothelial cells,^43^ and with an earlier report have confirmed CREBBP/EP300 interactions with SMAD proteins in transfected 293T cells.^8^ Moreover, Narita et al. demonstrated that A-485-mediated inhibition of CREBBP/EP300 did not alter the chromatin occupancy, yet abolished enhancer activation, demonstrating that productive gene activation requires co-activator recruitment and acetyltransferase activity downstream of DNA occupancy.^44^ At present, it is not clear whether acetylation of histones or of other proteins is critical for TGF-β mediated activation, which would be a subject for further studies.

In conclusion, our work shows that our set of SMAD4 MH2 mutants is defective in the regulation of TGF-β responsive genes despite their efficient genome binding properties. The SMAD4 mutants display a reduced interaction with CREBBP/EP300 co-activators providing a molecular mechanism for the defective activation of TGF-β-induced genes by SMAD4 MH2 mutants.

## Authors contribution

**Md Saiful Islam:** Conceptualization; data curation; formal analysis; investigation; methodology; writing - original draft; writing - review and editing.

**Sheikh Nizamuddin:** Data curation; formal analysis; writing - original draft; writing - review and editing.

**Timothy En Haw Chan:** Investigation; methodology; writing - review and editing.

**Omid Fotouhi:** Investigation; writing - review and editing.

**Stefanie Koidl:** Investigation; writing - review and editing.

**H.T. Marc Timmers:** Conceptualization; funding acquisition; methodology; project administration; supervision; writing - review and editing.

## Acknowledgements

We thank the Lighthouse core facility of the Uniklinik Freiburg, the Genomics and Proteomics core facility of DKFZ, Heidelberg and the Deep Sequencing Facility Max Planck Institute of Immunobiology and Epigenetics, Freiburg. We are grateful to the members of Timmers lab for their helpful discussions and suggestions regarding this work. We acknowledge the use of BioRender app for the generation of Figures. This work was financially supported by the Deutsche Forschungsgemeinschaft (DFG) via SFB850, project B9.

