## Supplementary Data for "SMAD4 MH2 Mutations Disrupt CREBBP/EP300 Recruitment and TGF-β-Induced Transcription in Colorectal Cancer"

**Supplementary Tables**

Table S1: Enrichment of the SMAD4 consensus binding motif for WT and SMAD4 mutants across TGF-β treatment time points.

Table S2. Differentially enriched SMAD4 peaks upon TGF-β treatment in WT and SMAD4 mutants cells.

**Supplementary Figures**

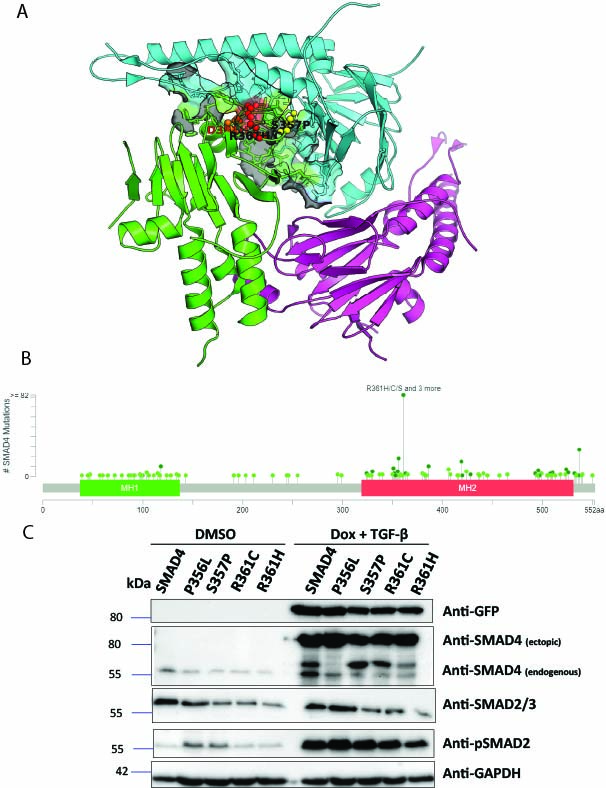

**Figure S1:** A) Distribution of SMAD4 MH2 domain missense mutations. The structure of SMAD4 is shown as a heterotrimeric complex of SAMD4 with R-SMADs (SMAD2, SMAD3) and the 3D structure were visualized using PDB file via PyMOL 3.1. Orange, red and yellow color stand for D351H, S357P, R361H/C mutants of SMAD4 protein. B) Mutational landscape of *SMAD4* gene mutations in CRC. A green dot indicates the position of missense (putative driver mutation), dark green dots stand for missense (passenger mutation). Y-axis denotes number of mutations and runs to 82 (cbioportal.org), C) GFP-tagged SMAD4 wt and mutant protein expression. RPE-1 cells expressing GFP-tagged SMAD4 were generated using doxycycline inducible FRT-based system. Cells were treated overnight with 1 µg/ml doxycycline.

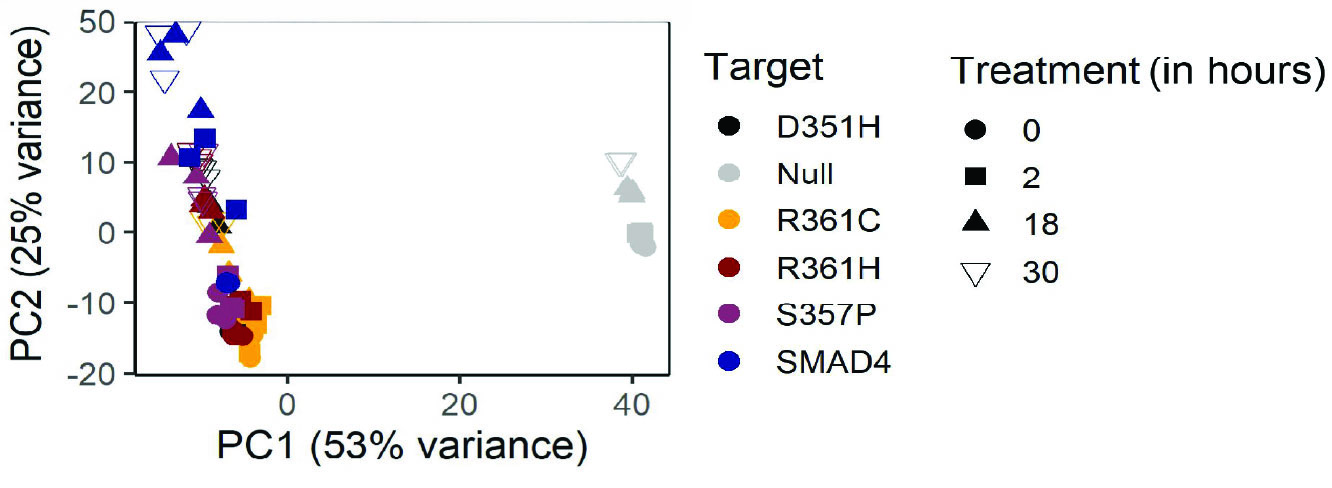

**Figure S2:** PCA analysis of RNA-seq data using the top differentially expressed 500 genes in SMAD4 wt, and mutants.

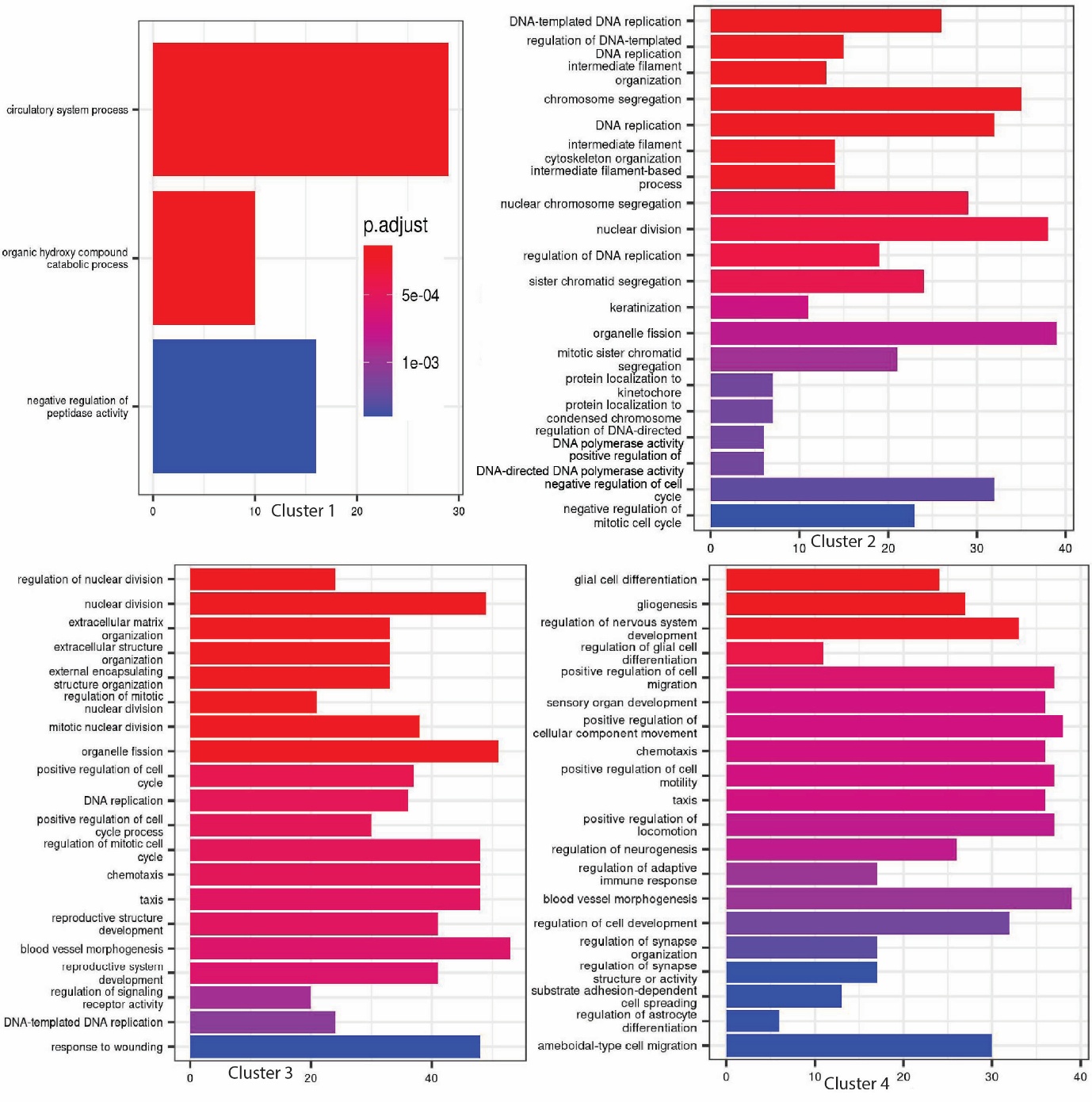

**Figure S3:** Biological pathway analysis of cluster 1, 2, 3, and 4 genes from the time course RNA-seq experiment of Fig. 2.

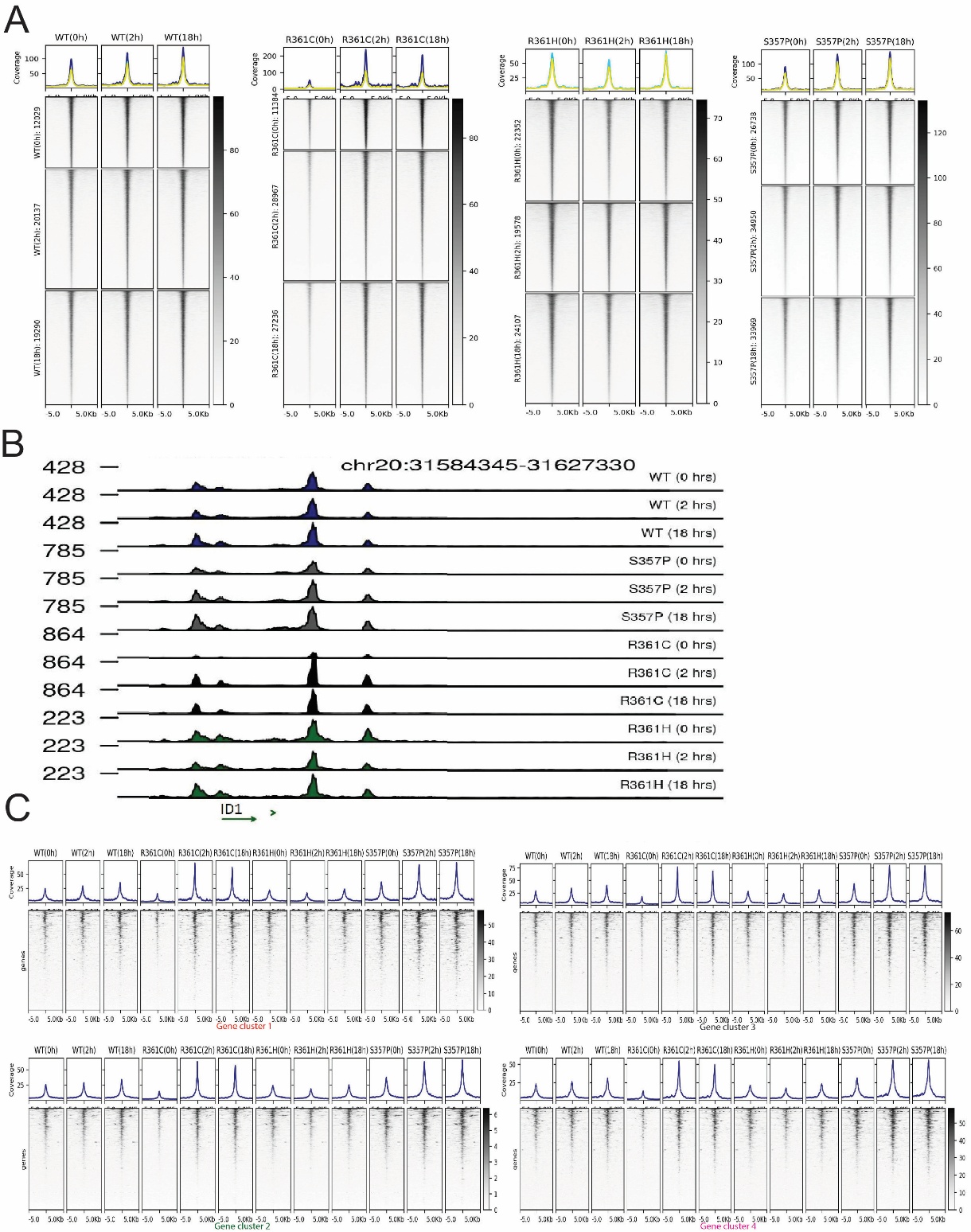

**Figure S4:** A) Heat maps representing SMAD4 wt and mutant binding after TGF-β treatment, B) Coverage track for SMAD4 binding site at *ID1* gene. C) Heat maps for wt and mutant SMAD4 binding at the promoter regions of RNA-seq cluster genes.

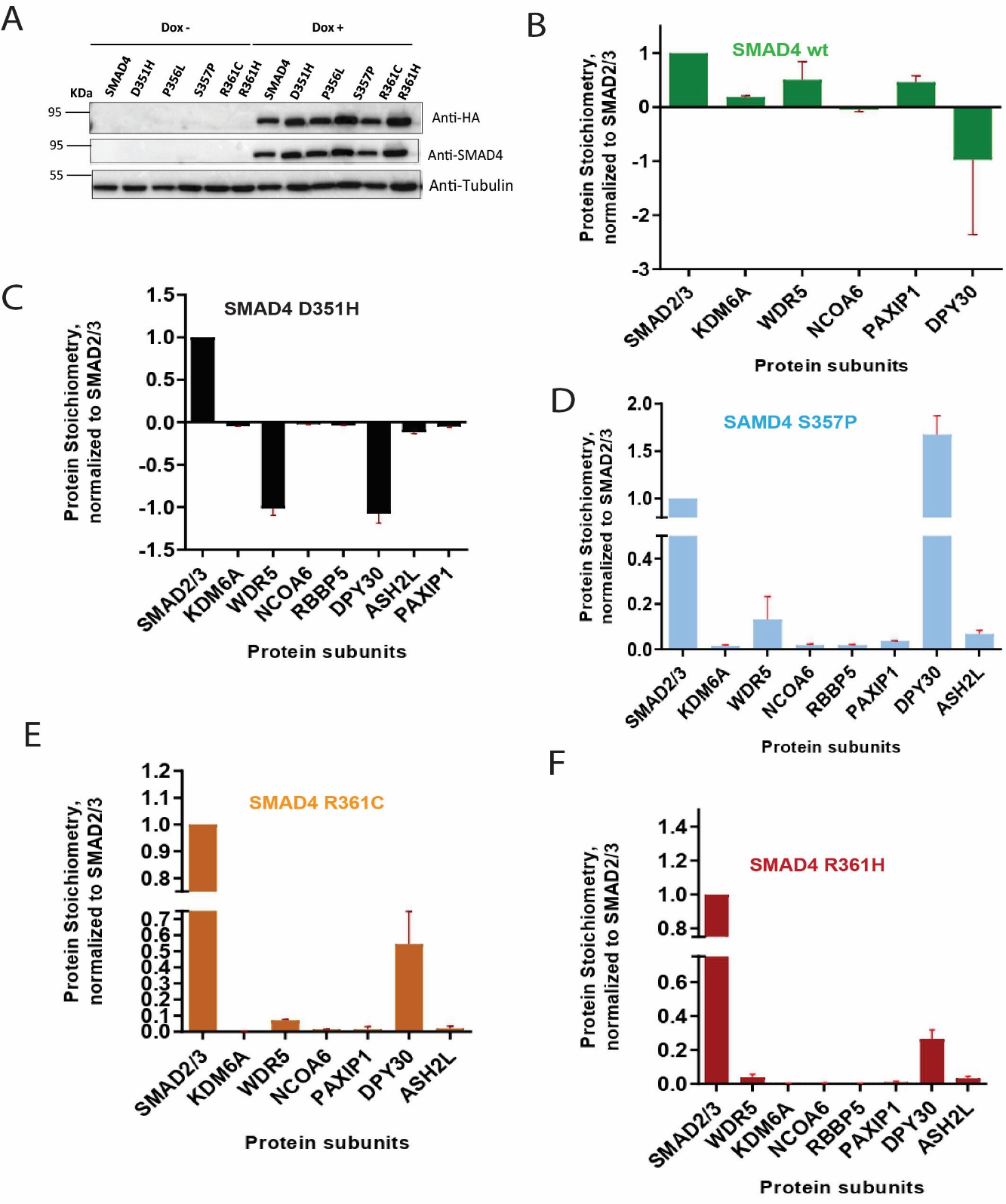

**Figure S5:** A) Immunoblot analysis of COLO205 cell lines expressing HA-miniTURBO fusions of wt and mutant SMAD4. (B-F) Stoichiometry analysis of MLL3/4 complex in wt and mutant SMAD4 cell lines.

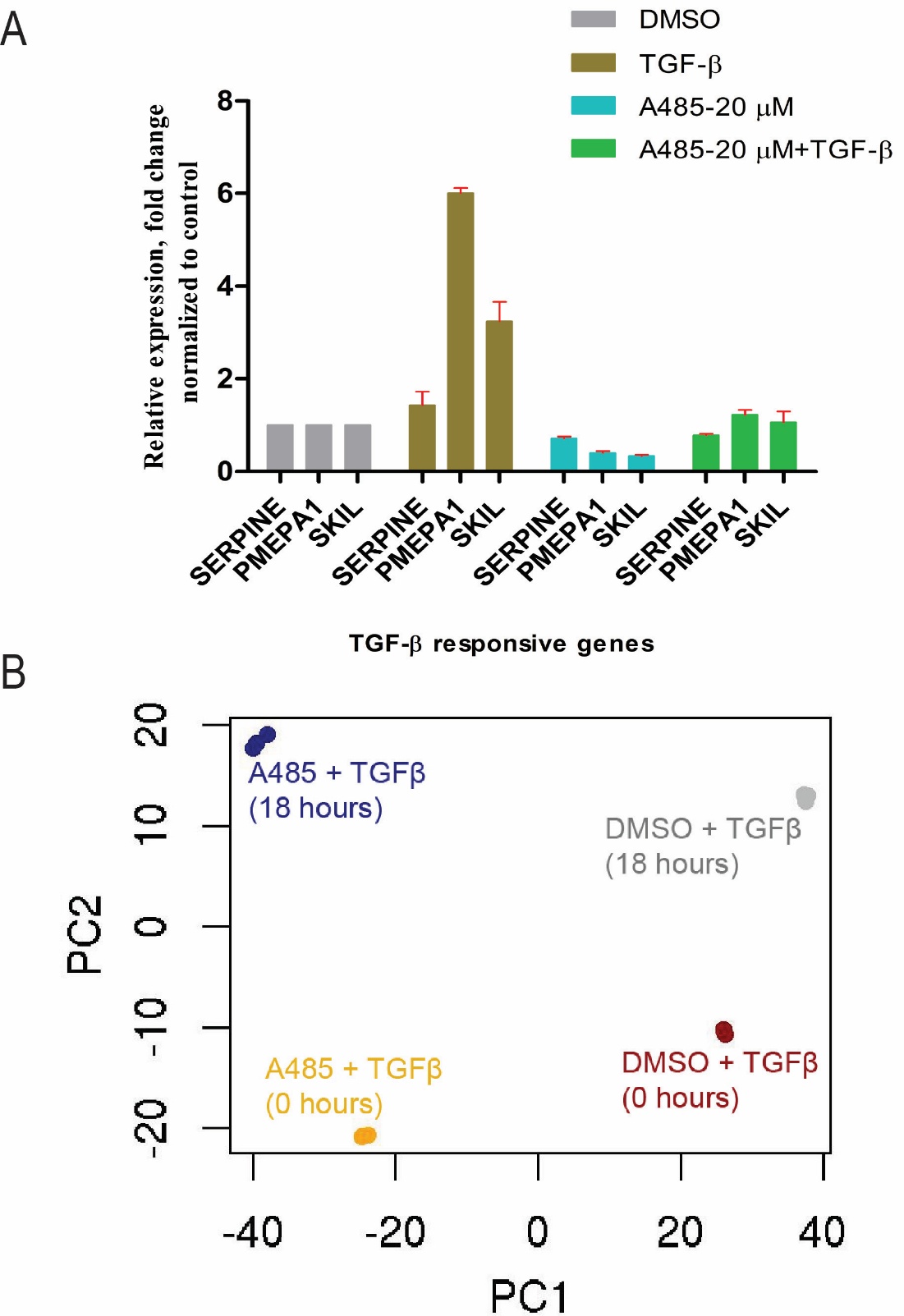

**Figure S6:** A) RT-qPCR analysis for TGF-β regulated gene expression. GFP-tagged SMAD4 wt cells were treated with A-485 for 4 hours followed by 10 ng/ml TGF-β or equal volume of DMSO treatment for 18 hours. Error bars depict the standard error of 2-4 biological replicates, B) PCA analysis of RNA-seq data using the top 500 DEGs.
