## Supplementary Information for "SMAD4 MH2 Mutations Disrupt CREBBP/EP300 Recruitment and TGF-β-Induced Transcription in Colorectal Cancer"

**List of antibodies for immunoblots**

| **Primary antibody** | **Dilution** | **Secondary antibody** | **Dilution** | **Company** |
| --- | --- | --- | --- | --- |
| GFP | 5% milk, 1:3000 | Mouse | 5% milk, 1:5000 | Santa Cruz sc #9996 |
| pSMAD2 | 5% milk+ 0.5% BSA, 1:1000 | Rabbit | 5% milk+ 0.5% BSA, 1:3000 | Cell Signalling #3108 |
| SMAD2/3 | 5% milk, 1:1000 | Mouse | 5% milk, 1:3000 | BD Biosciences #610842 |
| SMAD4 | 5% milk+ 0.5% BSA, 1:1000 | Rabbit | 5% milk+ 0.5% BSA, 1:3000 | Cell Signalling #9515S |
| HA | 5% milk, 1:3000 | Mouse | 5% milk, 1:5000 | 12CA5 ascites |
| alpha-Tubulin | 5% BSA, 1:1000 | Rabbit | 1X TBST, 1:2000 | Cell Signaling # 2144 |
| Vinculin | 5% milk, 1:1000 | Goat | 5% milk, 1:5000 | Santa Cruz sc #7649 |
| GAPDH | 5% milk, 1:1000 | Rabbit | 5% milk, 1:5000 | Cell Signaling #5174 |

**List of primers for qRT-PCR**

| **Primers Name** | **Forward** | **Reverse** |
| --- | --- | --- |
| *SERPINE* | AAGGCACCTCTGAGAACTTCA | CCCAGGACTAGGCAGGTG |
| *PMEPA1* | CTGTCTGCACGGTCCTTCAT | CCACAGGCATCCTTCTGAGG |
| *SKIL* | GAGGCTGAATATGCAGGACAG | CTTGCCTATCGGCCTCAG |
| *TBP* | TGCACAGGAGCCAAGAGTGAA | CACATCACAGCTCCCCACCA |
| *GAPDH* | TGCACCACCAACTGCTTAGC | GGCATGGACTGTGGTCATGAG |
| *HMBS* | GGCAATGCGGCTGCAA | GGGTACCCACGCGAATCAC |
| *HPRT1* | TGACACTGGCAAAACAATGCA | GGTCCTTTTCACCAGCAAGCT |
| *ACTB* | CACCATTGGCAATGAGCGGTTC | AGGTCTTTGCGGATGTCCACGT |
